# Longitudinal changes in the prevalence and intensity of soil-transmitted helminth infection following expanded community-wide mass drug administration in the delta region of Myanmar

**DOI:** 10.1101/341677

**Authors:** Julia C Dunn, Alison A Bettis, Nay Yee Wyine, Aye Moe Moe Lwin, Aung Tun, Nay Soe Maung, Roy M Anderson

## Abstract

Mass drug administration (MDA), targeted at school-aged children is the method recommended by the World Health Organization for the control of morbidity induced by soil-transmitted helminth (STH) infection in endemic countries. However, MDA does not prevent reinfection between treatment rounds. In countries with endemic infection, such as Myanmar, the MDA coverage, who is targeted, and rates of reinfection in given environmental and social settings will determine how effective mass drug treatment is in suppressing transmission in the long-term. In this paper, data from an epidemiology study on STH, conducted between June 2015 and June 2016 in the delta region of Myanmar, are analysed to determine the risks of STH infection in the whole community over a year which included two MDA rounds. Risk ratios (RRs) for the four-month reinfection period were below one, whereas RRs for the six-month reinfection period were above one, indicating that more people were infected after six months of exposure post-MDA. Evidence of predisposition, as measured by the Kendall Tau-b statistic, was found for all STH species and across all age groups. This study demonstrates that a six-month gap between MDA in these communities is enough time for STH infection to return to pre-MDA levels and that the same individuals are being consistently infected between MDA rounds.

**Author summary:** Mass drug administration (MDA), treating either whole communities or targeted groups without a prior diagnosis, is used as a control strategy for many neglected tropical diseases, including soil-transmitted helminth (STH) infection. MDA takes place at set intervals, aiming to reduce morbidity caused by the target disease and potentially interrupt transmission. In this study we measure STH infection in two villages in the delta region of Myanmar over the course of a year, both before and after MDA rounds, to quantify the effect of treatment on infection and to identify groups with persistent infections. We found that whilst overall prevalence of STH infection decreased over the year, intensity of infection, measured by eggs per gram of faeces, did not significantly decrease. We also found evidence to suggest that particular people are predisposed to STH infection. This is possibly due to non-compliance to MDA, or behavioural and social factors. The findings presented here will provide evidence to support continuing Myanmar’s MDA programme for STH control and using accurate diagnostics to identify and target “predisposed” people for sustained treatment.

## Introduction

Soil-transmitted helminth infections (STHs) are classified by the World Health Organization (WHO) as neglected tropical diseases (NTDs). Approximately 1.4 billion people worldwide are estimated to be infected with at least one of the main STHs (*Ascaris lumbricoides*, *Trichuris trichiura*, *Ancylostoma duodenale*, *Necator americanus*) (1). Endemic countries carry out mass drug administration (MDA) campaigns to control STH infections with the goals of reducing STH prevalence and intensity of infection to a level where there is a low risk of morbidity in children (2,3). The WHO recommends that MDA is carried out annually or biannually, targeting school-aged children (SAC, 5-14 years old) as they are at the highest risk of morbidity (3,4). In 2017, this guideline was updated to include treatment of young children (12 to 23 months old), preschool-aged children (pre-SAC, 2-4 year olds), adolescent girls (10-19 year olds) and women of reproductive age (WRA, 15-45 year olds) (3). Myanmar, in Southeast Asia, has conducted pre-SAC and SAC targeted MDA with albendazole every August since 2003 and reported high coverage levels. For example, national coverage of SAC was reported as 97.49% in 2016 (5,6). However, contrary to the usual WHO definitions of SAC (5-14 years old (4)), the STH MDA campaigns in Myanmar only treats pre-SAC and primary school children (5-9 year olds) (5). A community-wide MDA with diethylcarbamazine citrate (DEC) and albendazole is carried out by the Global Programme to Eliminate Lymphatic Filariasis (GPELF) in December or January of each year (7). Therefore, children 2-9 years old get treated with albendazole biannually whereas anyone 10 years and over get treated with albendazole annually. Whilst intensity of STH has decreased to low levels since MDA began, prevalence is still too high to consider halting the MDA programme (5,8,9).

The purpose of preventive chemotherapy is to clear STH infections from humans, but it does not prevent reinfection between MDA rounds since gut dwelling helminth parasites do not trigger strong acquired immunity in the human host (10). Research into drug efficacy also suggests that a single dose of albendazole, as given in most countries’ MDA programmes, will not completely clear intestinal helminth parasites, especially *T. trichiura* infections (11–14). Therefore, as well as individuals gaining new infections after MDA, they may also be harbouring old infections not killed by previous treatment. Reinfection, or the change in STH prevalence and intensity over time, depends on multiple factors: the efficacy of the anthelminthics and coverage of MDA to effectively clear STH infections from all infected individuals in the population, the level of environmental contamination with eggs and larvae and an individuals’ exposure to environmental contamination (behavioural and social factors) (15).

Those who are consistently reinfected with STH after clearing their infections with treatment are considered to be “predisposed” to infection (16). Research is ongoing to determine what the underlying factors of predisposition are. They are likely to be a combination of genetic, immunological, environmental and behavioural factors (17–19). Predisposition is usually defined as individuals who consistently reacquire infection to the same intensity class of infection (i.e. low, medium, high) as prior to treatment (20). However, in low STH intensity populations, such as those that have undergone multiple years of MDA, the definition of predisposition could be widened to include all those who consistently harbour positive STH infections between MDA rounds. Being able to identify the groups of people that are consistently infected, despite MDA, would be highly beneficial to STH control programmes that are in the final years of MDA and are targeting the interruption of transmission (21,22). A change in treatment policy may be desirable to target those predisposed to infection if they can easily be identified.

In this paper we describe and analyse the longitudinal infection profiles of participants in an epidemiology study of STH in Myanmar that was conducted between June 2015 and June 2016. The aim of this analysis is to determine how infection fluctuates over the course of a year under the influence of the regular government-led MDA, and expanded community-based MDA, both of which aim to significantly reduce STH prevalence and intensity. We also investigate if there is evidence for predisposition to infection in the study sample when stratified by a variety of confounding variables including age and sex.

## Methods

### Study sites and design

Data were collected in an STH epidemiological study that has been detailed in a previous publication (8). The study received full ethical approval from the Imperial College Research Ethics Committee, Imperial College London, UK and the Department of Medical Research Ethics Review Committee, Ministry of Health and Sports, Myanmar. Data collection took place between June 2015 and June 2016 in Udo village, Taikkyi township, Yangon Region and Kyee Kan Theik village, Nyaung Don township, Ayeyarwaddy Region. Further details on the study sites, including environmental and socioeconomic information, are provided in the previous publication (8). In June 2015, a demographic survey and census were completed in the two study villages. Participants for the study were randomly selected by household. Participants completed questionnaires collecting data on participants’ socioeconomic status, household structure and access to water, sanitation and hygiene (WaSH) facilities. The study comprised three parasitology surveys in August 2015 (first survey – S1), December 2015 (second survey – S2) and June 2016 (third survey – S3) (Figure 1). Stool samples were collected from the participants in each parasitology survey and were assessed for STH infection using the Kato-Katz method (23). The participants and their stool samples were assigned unique identification (ID) codes to maintain confidentiality and to link results over all surveys. Treatment efficacy was measured by collecting stool samples from a sub-sample of participants two weeks after the first survey (August 2015), post-MDA, and assessing them for STH infection by Kato-Katz.

**Fig 1:**
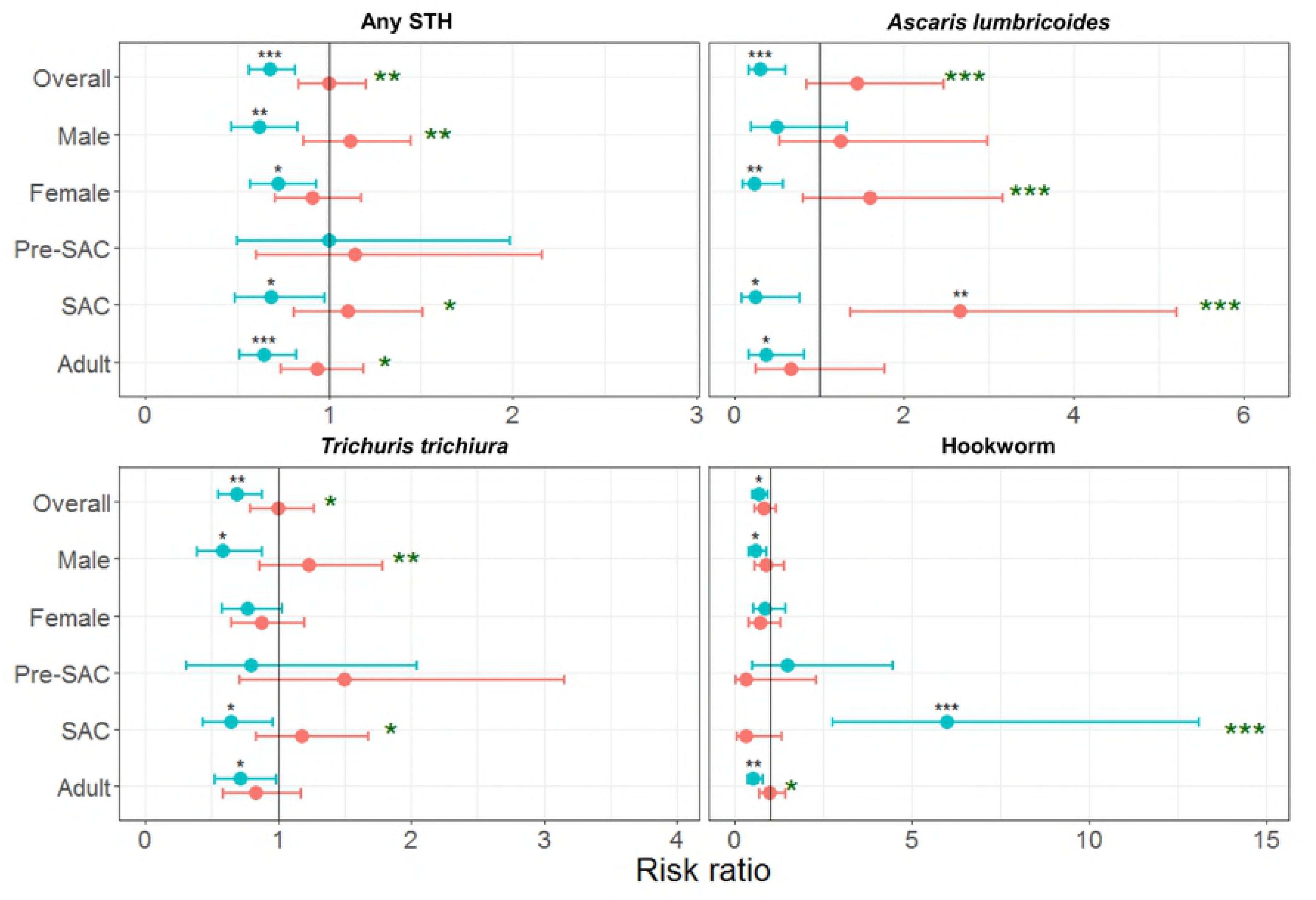
Flow diagram of data collection and study methods. *DEC* = *diethylcarbamazine citrate. Reproduced with permission from the supplementary information of Dunn et al. (2017)* (8).

### Data

Data for the following analyses were from all participants who had a recorded Kato-Katz result from all three surveys. Overall, 523 participants from 211 households had the requisite data. Data from both villages were merged and analysed as one dataset. Egg counts, measured by the Kato-Katz method, were multiplied by 24 to give eggs per gram of faeces (EPG) (24). All data were anonymised and assigned a unique ID code to ensure that data could be linked over the course of the study.

### Statistical analysis

RStudio (R version 3.0.1, Vienna, Austria) was used for the following statistical analyses and to create the figures. Participants were grouped into age groups as defined by the WHO: preschool-aged children (pre-SAC) are 2-4 year olds, school-aged children (SAC) are 5-14 year olds and adults are 15+ year olds (4). Exact confidence intervals (95% two-sided) for mean prevalence were calculated using the Clopper-Pearson method (25). Mean EPG adjusted percentiles (bias-corrected and accelerated - BCa) were calculated using bootstrapping methodology with the *“boot”* package. The significance test for the differences between risk ratios was derived from a formula published by Altman and Bland, 2003 (26). The WHO recommended intensity cut-offs were used to group individual EPG into low, medium and high intensity infections (2). To assess the differences in prevalence and intensity of infection over the surveys we used generalised linear models (GLMs) with a logit link (for binomial prevalence outcome) or a log link (for negative binomial intensity outcome) and the significance level was set at *P* ≤ 0.05. Kendall’s Tau-b values were calculated to assess predisposition to infection, adjusting for tied ranks, and the significance level set at *P* ≤ 0.05.

The study took place over the course of a year. Therefore, all participants will have aged one year during the study. Whilst there is a well-established relationship between age and STH infection, for simplicity we maintained the recorded ages for all participants at the age recorded in the first survey. This is assuming that age-related exposure did not drastically change over the course of a year. Also, we have maintained the usual WHO definition of SAC (5-14 years old), despite the fact that there is a different treatment frequency for 5-9 and 10-14 year olds in Myanmar. We have done this to align with how the WHO expects STH outcomes to be reported regarding infection prevalence and intensity by age grouping.

## Results

### Response to treatment

Out of the 523 participants with full Kato-Katz data, 60 (11.47%) were assessed for response to treatment (i.e. treatment efficacy). Only 15 of the 60 participants had positive STH infections in the pre-MDA survey and these individuals were therefore used for assessing STH clearance. Due to the small sample size, response to treatment cannot be accurately quantified and we cannot guarantee that all positive infections were cleared after MDA. Therefore, “reinfection” in the case of our analysis does not necessarily refer to new infections picked up between MDA rounds, but rather changes in the number and proportion of positive infections between surveys.

### Reinfection – prevalence

Over the year (S1 to S3), prevalence of any STH fell by 8.99% and the reduction was statistically significant (*P* < 0.001). The reductions in prevalence of each STH separately, were also statistically significant (*P* < 0.05). Prevalence of infection with at least one STH fell between S1 and S2 and was maintained between S2 and S3 (Table 1). This was also true for *T. trichiura* prevalence where the efficacy of albendazole to clear infection is much less that for *A. lumbricoides* and hookworm. *A. lumbricoides* infection fell and then rose slightly. Hookworm prevalence decreased over the course of the three surveys.

**Table 1:**
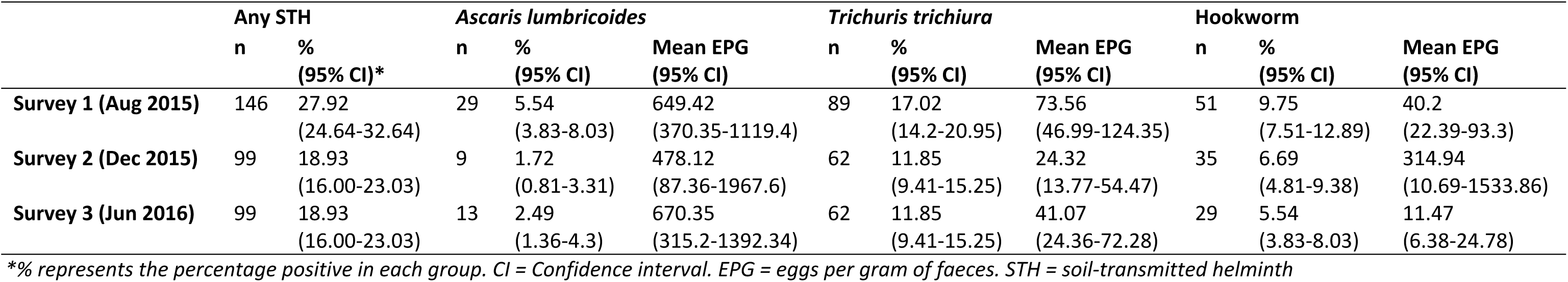
Number of positive individuals (n), prevalence (%) and infection intensity of each soil-transmitted helminth species (overall n=523)

Risk ratios (RRs) for reinfection differed between STH species and reinfection period. RRs for the four-month reinfection period (S1 to S2) were mostly below one, indicating that infection decreased between these surveys (Figure 2, S1 Table). Most RRs for the six-month reinfection period (S2 to S3) were above one but the CIs for all six-month RRs cross one, except for *A. lumbricoides* infection in SAC, indicating non-significance. There were few deviations from the overall patterns of RRs when stratified by sex and age group. Six-month RRs were significantly higher than four-month RRs within the sample for all species except hookworm, and were significantly higher within stratifications, especially SAC (any STH, *A. lumbricoides* and *T. trichiura*), adults (any STH and hookworm) and males (any STH and *T. trichiura*). However, inversely, the four-month RR for hookworm infection in SAC was significantly higher than the six-month RR.

**Fig 2:**
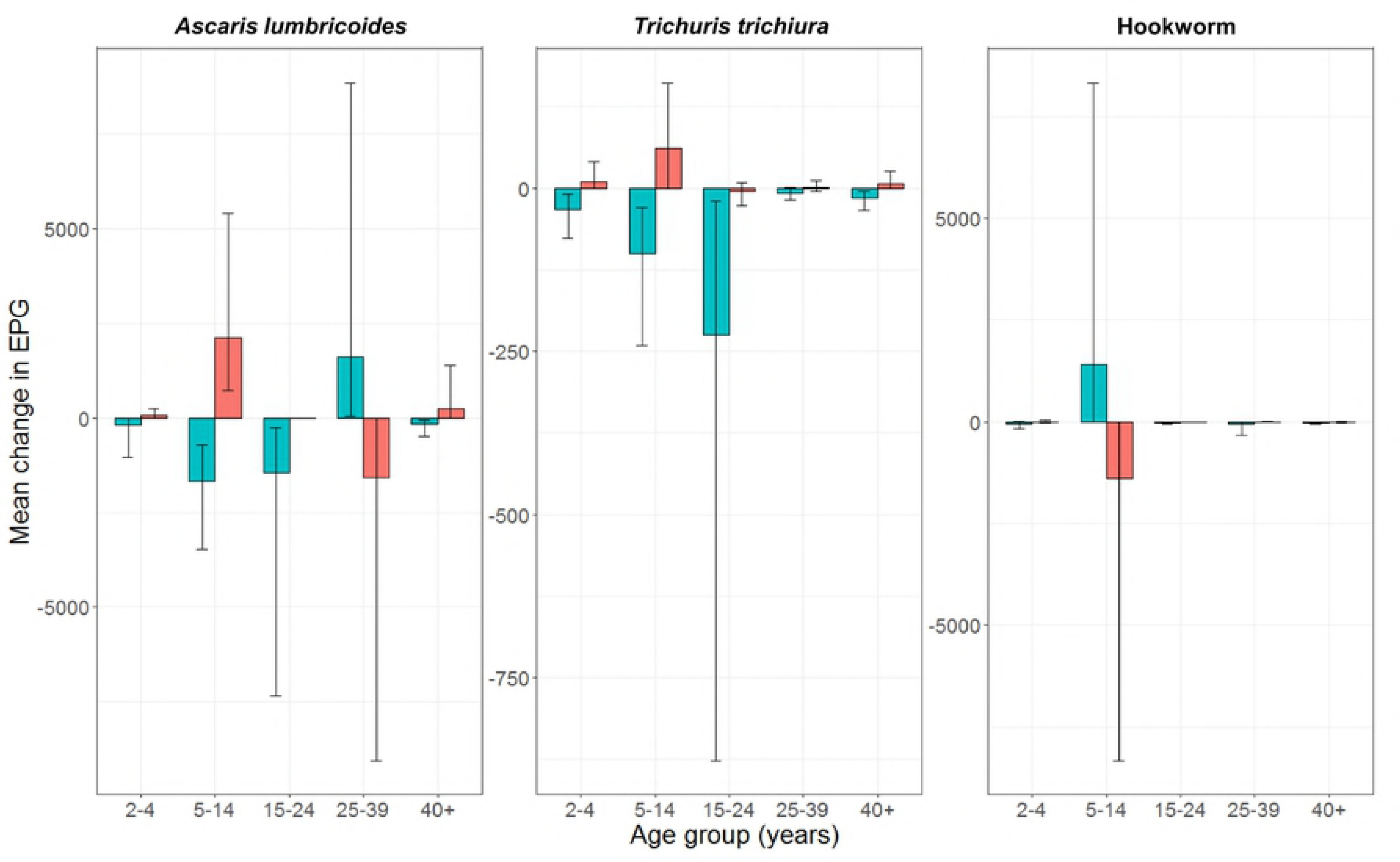
Risk ratios of STH prevalence between surveys. Blue = 4 months reinfection (survey 1 to survey 2). Red = 6 months reinfection (survey 2 to survey 3). * P ≤ 0.05, ** P ≤ 0.01, *** P ≤ 0.001 – black asterisks represent statistical significance of each risk ratio, green asterisks represent statistical difference between risk ratios for each group. Horizontal lines represent 95% confidence intervals. Pre-SAC = preschool-aged children (2-4 years old), SAC = school-aged children (5-14 years old), Adult = 15+ years old. No positive Ascaris lumbricoides infections were found in pre-SAC for all surveys, therefore no points are presented.

### Reinfection – intensity

Over the year (S1 to S3), EPG decreased for *T. trichiura* and hookworm, but rose for *A. lumbricoides* (Table 1). However, the only statistically significant difference was for the decrease in hookworm EPG from 40.20 to 11.47 (*P* < 0.05). A majority of the positive STH infections in all three surveys were low intensity infections. Over all the surveys and species infections, of the 98 instances that infections moved from negative in the prior survey to positive, 88 (89.80%) moved to low intensity, seven (7.14%) to medium intensity and three (3.06%) to high intensity (Table 2). Of the 95 instances where individuals were reinfected after treatment to the same intensity group as recorded in the prior survey, 87 (91.58%) were low to low intensity, eight (8.42%) were medium to medium intensity, and none were high to high intensity.

**Table 2:** Percentage and number (n) of individual infections between infection and intensity groups (overall n = 523)

|  | *Ascaris lumbricoides* |  | *Trichuris trichiura* |  | Hookworm |  |
| --- | --- | --- | --- | --- | --- | --- |
|  | S1-S2 | S2-S3 | S1-S2 | S2-S3 | S1-S2 | S2-S3 |
| Never infected (neg-neg) | 93.50 (489) | 96.75 (506) | 79.16 (414) | 82.22 (430) | 86.81 (454) | 90.25 (472) |
| Cured (pos-neg) | 4.78 (25) | 0.76 (4) | 8.99 (47) | 5.93 (31) | 6.50 (34) | 4.21 (22) |
| Negative to infected (neg-pos) | 0.96 (5) | 1.53 (8) | 3.82 (20) | 5.93 (31) | 3.44 (18) | 3.06 (16) |
| Decrease intensity group (pos-pos) | 0.38 (2) | 0 | 0.96 (5) | 0.38 (2) | 0.19 (1) | 0 |
| Same intensity group (pos-pos) | 0.19 (1) | 0.57 (3) | 6.88 (36) | 4.97 (26) | 3.06 (16) | 2.49 (13) |
| Increase intensity group (pos-pos) | 0.19 (1) | 0.38 (2) | 0.19 (1) | 0.57 (3) | 0 | 0 |

Pos = positive. Neg= negative. S1 = survey 1 (August 2015), S2 = survey 2 (December 2015), S3 = survey 3 (June 2016).

Overall mean EPG decreased from S1 to S2 but increased from S2 to S3 for *A. lumbricoides* and *T. trichiura*. Hookworm mean EPG increased from S1 to S2 and decreased from S2 to S3 (Table 1). When mean EPG change was stratified by age group (Figure 3, S2 Table) these patterns were reflected for all age groups except the 25-39 year olds for *A. lumbricoides*. The mean change in EPG was not homogenous between all age groups. There was minimal change in mean EPG for both *A. lumbricoides* and *T. trichiura* in the youngest and oldest age groups. For hookworm, the increase and decrease in mean EPG was driven by the change in 5-14 year olds.

**Fig 3:**
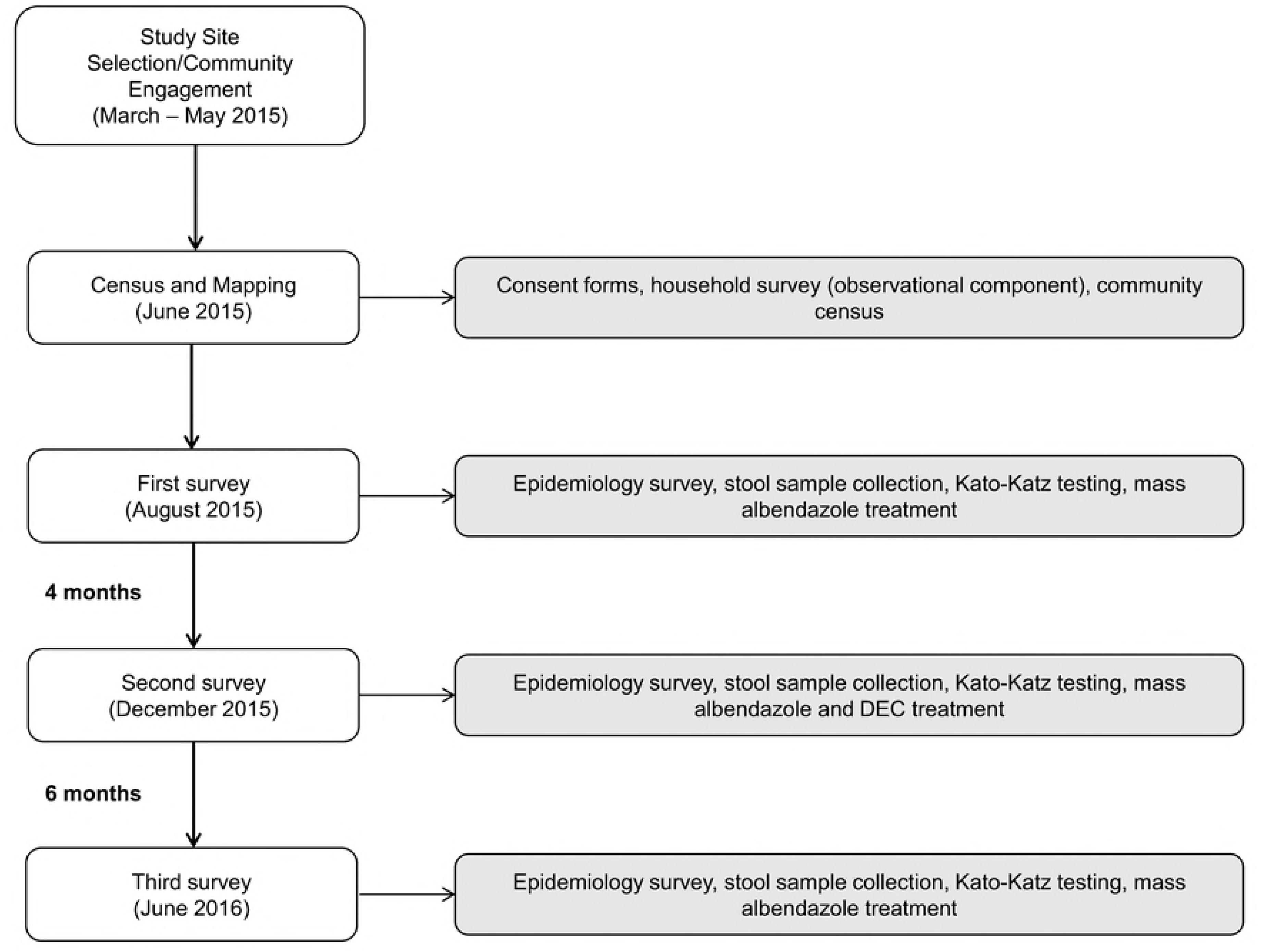
Mean change in eggs per gram of faeces (EPG) by age group. Red bars = 4 months reinfection (survey 1 to survey 2). Blue bars = 6 months reinfection (survey 2 to survey 3). Vertical lines represent 95% confidence intervals.

### Predisposition to STH infection (consistent infection)

A total of 38 (7.27%) participants had STH positive infections in all three surveys and 67 (12.81%) had positive infections for any two of the three surveys. Correlation coefficients (Kendall Tau-b) of individual participants’ egg count results between surveys were statistically significant for all species of STH (Table 3). Most of the correlations remained significant when stratified by sex or age group. Tau values ranged between zero and one, a higher value indicates stronger concordance (27). *Trichuris trichiura* egg counts had the strongest concordance between surveys especially for males, SAC and pre-SAC. The strongest concordance was for hookworm egg counts in pre-SAC between the first and second surveys. However, the Kendall’s Tau-b value may have been inflated due to the small number of pre-SAC infected with hookworm (two in survey one, three in survey two). Non-significant and low Tau values were calculated for *A. lumbricoides* infection in males and hookworm infection in SAC, denoting little evidence for predisposition in these groups.

**Table 3:** Kendall’s Tau-b correlation coefficients for individual participants’ egg counts between surveys

|  |  | *Ascaris lumbricoides* |  |  | *Trichuris trichiura* |  |  | Hookworm |  |  |
| --- | --- | --- | --- | --- | --- | --- | --- | --- | --- | --- |
|  |  | S1-S2 | S2-S3 | S1-S3 | S1-S2 | S2-S3 | S1-S3 | S1-S2 | S2-S3 | S1-S3 |
| Overall |  | 0.23*** | 0.45*** | 0.23*** | 0.50*** | 0.43*** | 0.48*** | 0.36*** | 0.36*** | 0.37*** |
| Sex | Male | 0.02 | 0.44*** | 0.14** | 0.54*** | 0.50*** | 0.58*** | 0.41*** | 0.43*** | 0.42*** |
|  | Female | 0.37*** | 0.45*** | 0.27*** | 0.47*** | 0.38*** | 0.40*** | 0.27*** | 0.27*** | 0.25*** |
| Age group | Pre-SAC | NA | NA | 0.03 | 0.62*** | 0.55*** | 0.41* | 0.81*** | 0.57*** | 0.67*** |
|  | SAC | 0.30* | 0.58*** | 0.23* | 0.62*** | 0.41*** | 0.56*** | 0.02 | 0.25* | 0.01 |
|  | Adults | 0.18*** | 0.40*** | 0.24*** | 0.40*** | 0.40*** | 0.38*** | 0.38*** | 0.37*** | 0.36*** |

\* $P \leq 0.05$ . \*\* $P \leq 0.01$ . \*\*\* $P \leq 0.001$ . S1 = survey 1 (August 2015). S2 = survey 2 (December 2015). S3 = survey 3 (June 2016). Kendall Tau-b values: Blue = 0.00-0.25, Green=0.26-0.50, Orange=0.51-0.75, Red=0.76-1.00, Grey=non-significant. Pre-SAC = preschool-aged children (2-4 years old). SAC = school-aged children (5-14 years old). Adults = 15+ years old. NA = comparison not possible (zero positive *Ascaris lumbricoides* infections in pre-SAC survey 2).

## Discussion

Myanmar has been conducting SAC-targeted MDA since 2003, and community-wide MDA in the delta region since 2013 (5,9). Whilst STH prevalence has dropped significantly since the initiation of MDA, the prevalence target set by WHO to discontinue MDA (under 1%) has not yet been reached in surveyed communities (8,28). Currently, there is no monitoring and evaluation (M&E) of STH in Myanmar, and no longitudinal studies have taken place since 1990 (29). It is therefore important for longitudinal M&E studies to take place in the country so that the long-term impact of MDA can be evaluated. The analyses in this study demonstrate that STH prevalence and intensity levels vary throughout the year. Overall prevalence of each STH was reduced following two community-wide MDA rounds (prevalence of any STH - 27.92-18.93%), and the intensities of infection (measured by EPG) of each STH, except *A. lumbricoides*, were reduced.

In this analysis, risk ratios were used to describe the patterns of infection over a four-month and six-month reinfection period. Four months post-MDA, the risk of STH infection was lower than in the preceding survey (RR=0.67, 95% CI 0.56-0.81). The only statistically significant six-month RR was for *A. lumbricoides* infection in SAC (RR=2.67, 95% CI 1.37-5.21). However, the six-month RRs were significantly higher than the four-month RRs for infection with all STH species except hookworm. Six-month RRs were also statistically significantly higher for SAC for infection with any STH, *A. lumbricoides* and *T. trichiura*, but were significantly lower for hookworm. The reinfection rates for STH species and the rapidity of bounce back to pre-treatment levels, are fast for *A. lumbricoides* and *T. trichiura*, and slower for hookworm. This is in part related to the fact that the dynamical timescale of each infection in its response to population disturbance as induced by MDA is directly related to adult worm life expectancy, which is around one year for *A. lumbricoides* and *T. trichiura* and about two years for hookworm (30). A study by Yap *et al*., conducted in China, measured rapid reinfection of *A. lumbricoides* (75.8% and 83.8% four and six months post-treatment, respectively), but not for *T. trichiura* and hookworm (31). Other studies have recorded STH reinfection after six months post-treatment, but not to above the pre-treatment prevalence levels (32,33). If we assume that the MDA rounds had cleared infection, then the data suggest that four months is not enough time for STH to reinfect individuals to the infection levels before that particular round of treatment, but six months may be enough time. It should also be noted that the third survey (June 2016) took place two months prior to the usual MDA timing (August) when only SAC are treated. As such, SAC will have a further two months for reinfection and adults will have a further six or seven months until the community-wide lymphatic filariasis (LF) MDA round. However, it is more likely that, due to sub-100% drug efficacy and non-compliance to treatment, some infections were retained after MDA, and six months was enough time for the surviving helminths to release sufficient eggs to trigger the acquisition of new *A. lumbricoides* infections in SAC. There is also the possibility of a seasonal effect on transmission and reinfection (34,35). The first and third surveys both took place during the dry season, whereas the second survey took place during the rainy season. Infective stage survival is known to be increased during rainy seasons (35,36).

Prevalence is used as a key STH epidemiological metric, but intensity of infection is more important as a determinant of morbidity (2). Whilst STH prevalence dropped significantly between the first and third surveys, the slight reductions in mean *T. trichiura* and hookworm EPG were not statistically significant. STH intensity at the beginning of the study was already at a very low level. Most participants with positive infections had low intensity infections. Prior work on the effect of long-term MDA programmes on STH have identified that substantial drops in STH prevalence and intensity in the first years of MDA may be followed by smaller reductions in subsequent years (37,38). For example, an eight year MDA programme in Burundi reported significant drops in prevalence in the first four years and no further decrease in the last four years (38). A monitoring survey in Kenya found that whilst prevalence of *T. trichiura* was significantly reduced after three years of MDA, mean EPG was not (39). The reasons for this may well be related to MDA coverage levels and individual compliance to treatment at multiple rounds of treatment (40). Few studies to date have recorded individual compliance to treatment but the persistence of low levels of prevalence may, in part, be due to persistent non-compliers to treatment (41).

In this study, the largest changes in EPG between surveys were found in the 5-14 years age group. Since the whole of the study sample was treated (not just SAC), any changes in EPG must be due to differences within the age groups (including compliance to treatment and behavioural factors) and not treatment efficacy. There was an increased risk of SAC to STH infection, especially *A. lumbricoides* and *T. trichiura*. Evidence of predisposition to STH infection has been found in several epidemiology studies (16,42,43) and the results of the Kendall’s Tau-b analysis indicates that predisposition to infection exists within the study sample. Stronger concordance between survey egg counts, and therefore stronger evidence for predisposition, was found in males and the younger age groups for *T. trichiura* and hookworm infection but only in females for *A. lumbricoides* infection. This is in agreement with Holland *et al.* (44), who found stronger evidence for *A. lumbricoides* predisposition in females, but in disagreement with Haswell-Elkins *et al.* (45) and Quinnell *et al.* (46), with females more predisposed to hookworm infection.

A limitation of this study, following on from the data collection study that preceded it, is the low sensitivity of the Kato-Katz technique as a diagnostic tool. It is highly possible that positive infections were missed due to its use (47). Another limitation is that, due to ethical reasons, the whole study sample (all ages) had to be treated during the MDA rounds that immediately followed the surveys, instead of the usual targeted ages (SAC only after the first and third surveys). The patterns of reinfection presented here therefore do not necessarily represent the patterns that will have occurred in previous years. This will have affected the results by increasing the drop in STH infection between surveys, potentially exaggerating sample-wide reinfection after MDA. Finally, as we could not confirm clearance of infection after MDA, the results may not be viewed as true “reinfection”. During data collection we attempted to ensure that treatment was taken via directly-observed therapy (DOT) where possible, but without data to confirm that infections were cleared we cannot assume this was always the case.

In this study the villages had already received over 10 years of MDA (both to treat STH and LF), yet low level STH infection persists. This may well be due to persistent non-compliers to multiple rounds of treatment who continue to release infective stages into the environment, as well as the perpetuation of those infective stages in the environment without improved WaSH to prevent it (48,49). The key epidemiological observation in this study is the low-level persistence of infection despite frequent community-based MDA, and the strong evidence for predisposition. In the long term, if diagnosis can be made more precise with new tools such as qPCR and the costs of such tests be greatly reduced, then future STH control may need to be based on targeted treatment to those predisposed to infection in order to eliminate transmission (50,51).

## Acknowledgements

We thank the Myanmar Ministry of Health and Sports for their continued support and advice. We thank Tom Churcher, Maria-Gloria Basáñez and Rachel Pullan for their comments on earlier drafts of this manuscript. We thank James Truscott for advice concerning data analysis and visualisation.

## Acronyms and abbreviations

Bca: Bias-corrected and accelerated
CI: Confidence interval
DOT: Directly observed therapy
EPG: Eggs per gram of faeces
GLM: Generalised linear model
GPELF: Global Programme to Eliminate Lymphatic Filariasis
ID: Identification
LF: Lymphatic filariasis
MDA: Mass drug administration
NTD: Neglected tropical disease
PCA: Principal components analysis
pre-SAC: Preschool-aged children
RR: Risk ratio
S1: Survey 1
S2: Survey 2
S3: Survey 3
SAC: School-aged children
SES: Socioeconomic status
STH: Soil-transmitted helminth
WaSH: Water, sanitation and hygiene
WHO: World Health Organization
WRA: Women of reproductive age

## Supplementary information

S1 Table: Risk ratios of STH infection between surveys

S2 Table: Mean change in eggs per gram of faeces (EPG) between surveys

S3: STROBE Checklist

## References

1. Pullan RL, Smith JL, Jasrasaria R, Brooker, SJ. Global numbers of infection and disease burden of soil transmitted helminth infections in 2010. Parasit Vectors. 2014;7(1).

2. WHO. Eliminating soil-transmitted helminthiases as a public health problem in children: progress report 2001-2010 and strategic plan 2011-2020. Geneva, Switzerland: World Health Organization; 2012.

3. WHO. Guideline: Preventive chemotherapy to control soil-transmitted helminth infections in at-risk population groups. WHO. Geneva: World Health Organization; 2017.

4. WHO. Helminth control in school-age children - Second Edition. Geneva, Switzerland: World Health Organization; 2012.

5. Tun A, Myat SM, Gabrielli AF, Montresor A. Control of soil-transmitted helminthiasis in Myanmar: results of 7 years of deworming. Trop Med Int Heal. 2013;18(8):1017–20.

6. WHO. PCT databank: soil-transmitted helminthiases [Internet]. Geneva, Switzerland: World Health Organization; 2017 [cited 2017 Jan 4]. Available from: http://www.who.int/neglected_diseases/preventive_chemotherapy/sth/en/

7. Padmasiri EA, Montresor A, Biswas G, de Silva NR. Controlling lymphatic filariasis and soil-transmitted helminthiasis together in South Asia: opportunities and challenges. Trans R Soc Trop Med Hyg. 2006;100(9):807–10.

8. Dunn JC, Bettis AA, Wyine NY, Lwin AMM, Lwin ST, Su KK, et al. A cross-sectional survey of soil-transmitted helminthiases in two Myanmar villages receiving mass drug administration: epidemiology of infection with a focus on adults. Parasit Vectors. 2017;10(374).

9. Montresor A, Zin TT, Padmasiri E, Allen H, Savioli L. Soil-transmitted helminthiasis in Myanmar and approximate costs for countrywide control. Trop Med Int Heal. 2004;9(9):1012–5.

10. Grencis RK. Immunity to helminths: resistance, regulation, and susceptibility to gastrointestinal nematodes. Annu Rev Immunol. United States; 2015;33:201–25.

11. Keiser J, Utzinger J. Efficacy of current drugs against soil-transmitted helminth infections. J Am Med Assoc. 2008;299(16):1937–48.

12. Belew S, Getachew M, Suleman S, Mohammed T, Deti H, D’Hondt M, et al. Assessment of efficacy and quality of two albendazole brands commonly used against soil-transmitted helminth infections in school children in Jimma Town, Ethiopia. PLoS Negl Trop Dis. 2015;9(9).

13. Sinniah B, Chew PI, Subramaniam K. A comparative trial of albendazole, mebendazole, pyrantel pamoate and oxantel pyrantel pamoate against soil-transmitted helminthiases in school children. Trop Biomed. 1990;7(2):129–34.

14. Stephenson LS, Latham MC, Kinoti SN, Kurz KM, Brighall H. Improvements in physical fitness of Kenyan schoolboys infected with hookworm, *Trichuris trichiura* and *Ascaris lumbricoides* following a single dose of albendazole. Trans R Soc Trop Med Hyg. 1990;84:277–82.

15. Cundill B, Alexander N, Bethony JM, Diemert D, Pullan RL, Brooker S. Rates and intensity of re-infection with human helminths after treatment and the influence of individual, household, and environmental factors in a Brazilian community. Parasitology. 2011;138(11):1406–16.

16. Schad GA, Anderson RM. Predisposition to hookworm infection in humans. Science. 1985 Jun;228(4707):1537–40.

17. Mangano VD, Modiano D. Host genetics and parasitic infections. Clin Microbiol Infect. England; 2014 Dec;20(12):1265–75.

18. Saltykova IV, Freydin MB, Ogorodova LM, Puzyrev VP. Genetic predisposition to helminthiases. Russ J Genet Appl Res. 2014;4(5):405–15.

19. Holland C V. Predisposition to ascariasis: patterns, mechanisms and implications. Parasitology. 2009 Oct;136(12):1537–47.

20. Walker M, Hall A, Basáñez MG. Individual predisposition, household clustering and risk factors for human infection with *Ascaris lumbricoides*: New epidemiological insights. PLoS Negl Trop Dis. 2011;5(4).

21. Truscott JE, Hollingsworth TD, Brooker SJ, Anderson RM. Can chemotherapy alone eliminate the transmission of soil transmitted helminths. Parasit Vectors. 2014;7(1):266.

22. Anderson R, Farrell S, Turner H, Walson J, Donnelly CA, Truscott J. Assessing the interruption of the transmission of human helminths with mass drug administration alone: optimizing the design of cluster randomized trials. Parasit Vectors. 2017;10(93).

23. Katz N, Chaves A, Pellegrino J. A simple device for quantitative stool thick-smear technique in Schistosomiasis mansoni. Rev Inst Med Trop Sao Paulo. 1972;14(6):397.

24. WHO. Bench aids for the diagnosis of intestinal parasites. Geneva, Switzerland; 1994.

25. Newcombe RG. Two-sided confidence intervals for the single proportion: comparison of seven methods. Stat Med. 1998;17:857–72.

26. Altman DG, Bland JM. Interaction revisited: the difference between two estimates. BMJ. 2003 Jan 25;326(7382):219.

27. Ng SK, Holden L, Sun J. Identifying comorbidity patterns of health conditions via cluster analysis of pairwise concordance statistics. Stat Med. John Wiley & Sons, Ltd; 2012 Nov 30;31(27):3393–405.

28. Htoon TT, Tun T, Oo KY, Thein W, Tin HH, Chai J-Y, et al. Status of infection with soil-transmitted helminths among primary school children in three selected townships of Yangon Division. Myanmar Heal Sci Res J. 2015;27(3).

29. Hlaing T, Saw T, Kyin M-L. Control of ascariasis through age-targeted chemotherapy: Impact of 6-monthly chemotherapeutic regimens. Bull World Health Organ. 1990;68(6):747–53.

30. Anderson RM, Medley GF. Community control of helminth infections of man by mass and selective chemotherapy. Parasitology. 1985;90(04):629–60.

31. Yap P, Du Z-W, Wu F-W, Jiang J-Y, Chen R, Zhou X-N, et al. Rapid re-infection with soil-transmitted helminths after triple-dose albendazole treatment of school-aged children in Yunnan, People’s Republic of China. Am J Trop Med Hyg. American Society of Tropical Medicine and Hygiene; 2013 Jul;89(1):23–31.

32. Sinniah B. Prevalence, treatment and re-infection of intestinal helminths among schoolchildren in Kuala Lumpur, Malaysia. Public Health. 1984;98(1):38–42.

33. Saathoff E, Olsen A, Kvalsvig JD, Appleton CC. Patterns of geohelminth infection, impact of albendazole treatment and re-infection after treatment in schoolchildren from rural KwaZulu-Natal/South-Africa. BMC Infect Dis. 2004 Dec 13;4(1):27.

34. Udonsi JK. Effectiveness of seasonal community-based mass-expulsion chemotherapy in the control of human hookworm infections in endemic communities. Public Health. 1985/09/01. 1985;99(5):295–301.

35. Davis EL, Danon L, Prada JM, Gunawardena SA, Truscott JE, Vlaminck J, et al. Seasonally timed treatment programs for Ascaris lumbricoides to increase impact—An investigation using mathematical models. PLoS Negl Trop Dis. 2018;12(1):e0006195.

36. Schulz S, Kroeger A. Soil contamination with *Ascaris lumbricoides* eggs as an indicator of environmental hygiene in urban areas of north-east Brazil. J Trop Med Hyg. 1992 Apr;95(2):95–103.

37. Supali T, Djuardi Y, Bradley M, Noordin R, Ruckert P, Fischer PU, et al. Impact of six rounds of mass drug administration on Brugian filariasis and soil-transmitted helminth infections in eastern Indonesia. PLoS Negl Trop Dis. 2013/12/19. 2013;7(12):e2586.

38. Ortu G, Assoum M, Wittmann U, Knowles S, Clements M, Ndayishimiye O, et al. The impact of an 8-year mass drug administration programme on prevalence, intensity and co-infections of soil-transmitted helminthiases in Burundi. Parasit Vectors. 2016 Dec 22;9(1):513.

39. Okoyo C, Nikolay B, Kihara J, Simiyu E, Garn J V, Freeman MC, et al. Monitoring the impact of a national school based deworming programme on soil-transmitted helminths in Kenya: the first three years, 2012 - 2014. Parasit Vectors. 2016 Jul;9(1):408.

40. Farrell S, Truscott J, Anderson RM. The importance of patient compliance in repeated rounds of mass drug administration (MDA) for the elimination of intestinal helminth transmission. Parasit Vectors. 2017;

41. Shuford K V., Turner HC, Anderson RM. Compliance with anthelmintic treatment in the neglected tropical diseases control programmes: a systematic review. Parasit Vectors. 2016 Dec 27;9(1):29.

42. Kepha S, Mwandawiro CS, Anderson RM, Pullan RL, Nuwaha F, Cano J, et al. Impact of single annual treatment and four-monthly treatment for hookworm and Ascaris lumbricoides, and factors associated with residual infection among Kenyan school children Multilingual abstracts. Infect Dis Poverty. 2017;6(30).

43. Wright JE, Werkman M, Dunn JC, Anderson RM. Current epidemiological evidence for predisposition to high or low intensity human helminth infection: a systematic review. Parasit Vectors. BioMed Central; 2018 Dec 31;11(1):65.

44. Holland C V, Asaolu SO, Crompton DW, Stoddart RC, Macdonald R, Torimiro SE. The epidemiology of Ascaris lumbricoides and other soil-transmitted helminths in primary school children from Ile-Ife, Nigeria. Parasitology. 1989 Oct;99 Pt 2(2):275–85.

45. Haswell-Elkins MR, Elkins DB, Manjula K, Michael E, Anderson RM. An investigation of hookworm infection and reinfection following mass anthelmintic treatment in the South Indian fishing community of Vairavankuppam. Parasitology. 1988;96(3):565–77.

46. Quinnell RJ, Slater AF, Tighe P, Walsh EA, Keymer AE, Pritchard DI. Reinfection with hookworm after chemotherapy in Papua New Guinea. Parasitology. 1993;106(4):379–85.

47. Easton A V, Oliveira RG, O’Connell EM, Kepha S, Mwandawiro CS, Njenga SM, et al. Multi-parallel qPCR provides increased sensitivity and diagnostic breadth for gastrointestinal parasites of humans: field-based inferences on the impact of mass deworming. Parasit Vectors. 2016 Jan 27;9(1):38.

48. Campbell SJ, Biritwum N-K, Woods G, Velleman Y, Fleming F, Stothard JR. Tailoring Water, Sanitation, and Hygiene (WASH) Targets for Soil-Transmitted Helminthiasis and Schistosomiasis Control. Trends Parasitol. 2017;1689.

49. Ziegelbauer K, Speich B, Mäusezahl D, Bos R, Keiser J, Utzinger J, et al. Effect of Sanitation on Soil-Transmitted Helminth Infection: Systematic Review and Meta-Analysis. PLoS Med. 2012 Jan 24;9(1):e1001162.

50. Easton A V., Oliveira RG, Walker M, O’Connell EM, Njenga SM, Mwandawiro CS, et al. Sources of variability in the measurement of Ascaris lumbricoides infection intensity by Kato-Katz and qPCR. Parasit Vectors. 2017 Dec 25;10:256.

51. Mejia R, Vicuña Y, Broncano N, Sandoval C, Vaca M, Chico M, et al. A Novel, Multi-Parallel, Real-Time Polymerase Chain Reaction Approach for Eight Gastrointestinal Parasites Provides Improved Diagnostic Capabilities to Resource-Limited At-Risk Populations. Am J Trop Med Hyg. The American Society of Tropical Medicine and Hygiene; 2013;88(6):1041–7.

